# Differential Methods for Assessing Sensitivity in Biological Models

**DOI:** 10.1101/2021.11.15.468697

**Authors:** Rachel Mester, Alfonso Landeros, Chris Rackauckas, Kenneth Lange

## Abstract

Differential sensitivity analysis is indispensable in fitting parameters, understanding uncertainty, and forecasting the results of both thought and lab experiments. Although there are many methods currently available for performing differential sensitivity analysis of biological models, it can be difficult to determine which method is best suited for a particular model. In this paper, we explain a variety of differential sensitivity methods and assess their value in some typical biological models. First, we explain the mathematical basis for three numerical methods: adjoint sensitivity analysis, complex-perturbation sensitivity analysis, and forward-mode sensitivity analysis. We then carry out four instructive case studies. (i) The CARRGO model for tumor-immune interaction highlights the additional information that differential sensitivity analysis provides beyond traditional naive sensitivity methods, (ii) the deterministic SIR model demonstrates the value of using second-order sensitivity in refining model predictions, (iii) the stochastic SIR model shows how differential sensitivity can be attacked in stochastic modeling, and (iv) a discrete birth-death-migration model illustrates how the complex-perturbation method of differential sensitivity can be generalized to a broader range of biological models. Finally, we compare the speed, accuracy, and ease of use of these methods. We find that forward-mode automatic differentiation has the quickest computation time, while the complex-perturbation method is the simplest to implement and the most generalizable.

**Author Summary:** Over the past few decades, mathematical modeling has become an indispensable tool in the biologist’s toolbox. From deterministic to stochastic to statistical models, computational modeling is ubiquitous in almost every field of biology. Because model parameter estimates are often noisy or depend on poorly understood interactions, it is crucial to examine how both quantitative and qualitative predictions change as parameter estimates change, especially as the number of parameters increase. Sensitivity analysis is the process of understanding how a model’s behavior depends on parameter values. Sensitivity analysis simultaneously quantifies prediction certainty and clarifies the underlying biological mechanisms that drive computational models. While sensitivity analysis is universally recognized to be an important step in modeling, it is often unclear how to best leverage the available differential sensitivity methods. In this manuscript we explain and compare various differential sensitivity methods in the hope that best practices will be widely adopted. In particular, we stress the relative advantages of existing software and their limitations. We also present a new numerical technique for computing differential sensitivity.

## 1 Introduction

In many mathematical models underlying parameters are poorly specified. This problem is particularly acute in biological and biomedical models. Model predictions can have profound implications for scientific understanding, further experimentation, and even public policy decisions. For instance, in an epidemic some model parameters can be tweaked by societal or scientific interventions to drive infection levels down. Differential sensitivity can inform medical judgement about the steps to take with the greatest impact at the least cost. Similar considerations apply in economic modeling. Finally, model fitting usually involves maximum likelihood or least squares criteria. These optimization techniques depend heavily on gradients and and Hessians with respect to parameters.

In any case it is imperative to know how sensitive model predictions are to changes in parameter values. Unfortunately, assessment of model sensitivity can be time consuming, computationally intensive, inaccurate, and simply confusing. Most models are nonlinear and resistant to exact mathematical analysis. Understanding their behavior is only approachable by solving differential equations or intensive and noisy simulations. Sensitivity analysis is often conducted over an entire bundle of neighboring parameters to capture interactions. If the parameter space is large or high-dimensional, it is often unclear how to choose representative points from this bundle. Faced with this dilemma, it is common for modelers to fall back on varying just one or two parameters at a time. Model predictions also often take the form of time trajectories. In this setting, sensitivity analysis is based on lower and upper trajectories bounding the behavior of the dynamical system.

The differential sensitivity of a model quantity is measured by its gradient with respect to the underlying parameters at their estimated values. Given the gradient of a function *f* (***β***), it is possible to approximate *f*(***β***) by the linear function *f*(***β***_0_) + ∇*f*(***β***_0_)^*t*^(***β*** – ***β***) for all ***β*** near ***β***_0_. The existing literature on differential sensitivity is summarized in the modern references [1, 2]. The Julia software DifferentialEquations.jl [3] makes sensitivity analysis fairly routine for many problems. Although the physical sciences have widely adopted the method of differential sensitivity [4, 5], the papers and software generally focus on a single sensitivity analysis method rather than a comparison of the various approaches. For example, PESTO [6] is a current Matlab toolbox for parameter estimation that uses adjoint sensitivities implemented as part of the CVODES method from SUNDIALS [7]. Should the continuous sensitivity equations be used, or would direct automatic differentiation of the solvers be more efficient on biological models? On the types of models biologists generally explore, would implicit parallelism within the sensitivity equations be beneficial, or would the overhead cost of thread spawning overrule any benefits? How close do simpler methods based on complex step differentiation get to these techniques? The purpose of the current paper is to explore these questions on a variety of models of interest to computational biologists.

In the current paper we also suggest a more accurate method of approximating gradients for models involving analytic functions without discontinuities, maxima, minima, absolute values or any other excursion outside the universe of analytic functions. By definition an analytic function can be expanded in a locally convergent power series around every point of its domain. In the sections immediately following, we summarize known theory, including the important adjoint method for computing the sensitivity of functions of solutions [4, 5]. Then we illustrate sensitivity analysis for a few deterministic models and a few stochastic models. Our exposition includes some straightforward Julia code that readers can adapt to their own sensitivity needs. These examples are followed by an evaluation of the accuracy and speed of the suggested numerical methods. The concluding discussion summarizes our experience, indicates limitations of the methods, and suggests new potential applications.

For the record, here are some notational conventions used throughout the paper. All functions that we differentiate have real or real vector arguments and real or real vector values. All vectors and matrices appear in boldface. The superscript ^*t*^ indicates a vector or matrix transpose. For a smooth real-valued function *f*(***x***), we write its gradient (column vector of partial derivatives) as ∇*f*(***x***) and its differential (row vector of partial derivatives) as *df*(***x***) = ∇*f*(***x***)^*t*^. If *g*(***x***) is vector-valued with *i*th component *g_i_*(***x***), then the differential (Jacobi matrix) *dg*(***x***) has *i*th row *dg_i_*(***x***). The chain rule is expressed as the equality *d*[*f* ∘ *g*(***x***)] = *df*[*g*(***x***)]*dg*(***x***) of differentials. The transpose (adjoint) form of the chain rule is ∇*f* ∘ *g*(***x***) = *dg*(***x***)^*t*^∇*f*[*g*(***x***)]. For a twice differentiable function, the second differential (Hessian matrix) *d*^2^*f*(***x***) = *d*∇*f*(***x***) is the differential of the gradient. Finally, *i* will denote 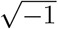.

## 2 Methods for Computing Sensitivity

The simplest dynamical models are governed by the linear constant coefficient differential equation 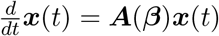 with solution 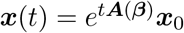. The directional derivative of the matrix exponential *e^**B**^* in the direction ***V*** can be represented by the integral

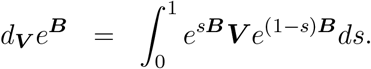

A proof of this fact appears in Example 3.2.2 of reference [8]. Setting ***B*** = *t**A***(***β***) and applying the chain rule leads to the partial derivative

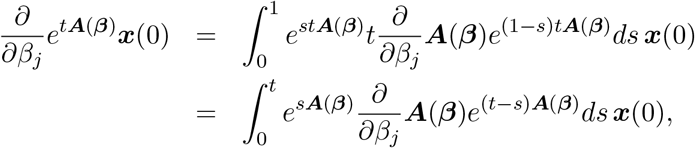

which can be laboriously evaluated by numerical integration. Simplification into a sum of exponentials is possible if ***A***(***β***) is uniformly diagonalizable across all ***β*** [9], but the details are messy.

In practice, it is simpler to differentiate the original ODE with respect to *β_j_*, interchange the order of differentiation, and numerically integrate the system

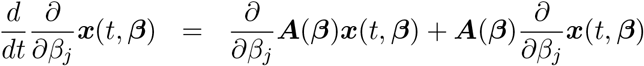

from 0 to some final value of *t*. The initial condition ***x***(0, ***β***) = ***x***(0) remains intact, and the new condition ∇***_β_x***(0, ***β***) = 0 is added. The second method has the advantage of giving the sensitivity along the entire trajectory. Another huge advantage is that the second method generalizes to nonlinear systems 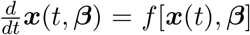 under the right circumstances. In particular the sensitivity of this more elaborate system can be evaluated by solving the differential equation

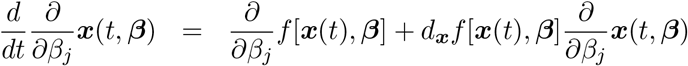

numerically. This sensitivity equation depends on knowing ***x***(*t*, ***β***). In practice, one solves the system

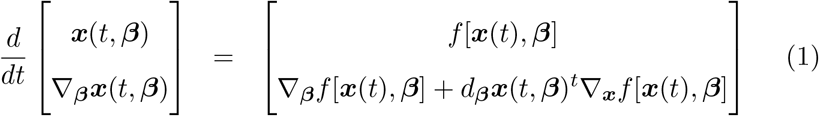

jointly, where *d**_β_x***[*t*, ***β***] is the Jacobi matrix of ***x***(*t*, ***β***) with respect to ***β***. This is commonly referred to as forward sensitivity analysis and is carried out by software suites such as DifferentialEquations.jl [3] and SUNDIALS CVODES [7]. We note in passing that a common implementation detail of sensitivity analysis is to base calculations on directional derivatives. Thus, the directional derivative

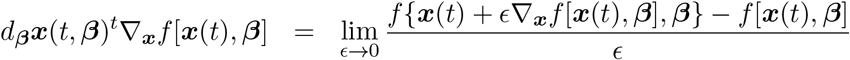

version of the chain rule allows one to evolve dynamical systems without ever computing full Jacobians.

The well-known adjoint method, another variant of the sensitivity method, is incorporated in the biological parameter estimation software PESTO through CVODES [7]. The adjoint method [1, 2] is defined directly on a function *g*[*x*(***β***), ***β***] of the solution of the ODE. The adjoint method first numerically solves the ODE system forward in time over [*t*_0_, *t_n_*], then solves the system

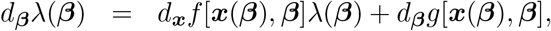

in reverse time, and finally computes derivatives via

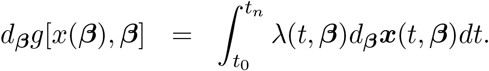

The second and third stages are commonly combined by appending the last equation to the set of ODEs being solved in reverse. This tactic achieves a lower computational complexity than other techniques, which require solving an *n*-dimensional ODE system *p* times for *p* parameters. In contrast, the adjoint method solves an *n*-dimensional ODE forwards and then solves an *n*-dimensional and a *p*-dimensional system in reverse, changing the computational complexity from 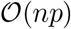 to 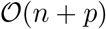. Whether such asymptotic cost advantages lead to more efficiency on practical models is precisely one of the points studied in this paper.

Alternatively, one can find the partial derivatives purely numerically. The obvious method here is to compute a slightly perturbed trajectory ***x***(*t*, ***β*** + Δ***v***) and form the forward differences

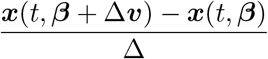

at all specified time points as approximations to the forward directional derivatives of ***x***(*t*, ***β***) in the direction ***v***. Choosing ***v*** to be unit vectors along each coordinate axis gives ordinary partial derivatives. The accuracy of this crude method suffers from round-off error in subtracting two nearly equal function values. These round-off errors are in addition to the usual errors committed in integrating the differential equation numerically. Round-off errors can be ameliorated by using central differences rather than forward differences, but this doubles computation time and still leaves the difficult choice of the stepsize Δ.

There is a far more accurate way when the function *f*[***x***, ***β***] defining the ODE is analytic in the parameter vector ***β*** [10]. This implies that the trajectory ***x***(*t*, ***β***) is also analytic in ***β***. For a real analytic function *g*(*β*) of a single variable *β*, the derivative approximation

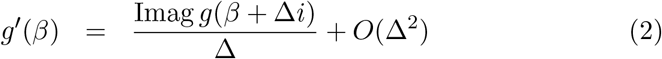

in the complex plane avoids roundoff and is highly accurate for Δ > 0 very small [11, 12]. Thus, in calculating a directional derivative of ***x***(*t*, ***β***), it suffices to (a) solve the governing ODE 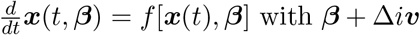 with ***β*** + Δ*i**v*** replacing ***β***, (b) take the imaginary part of the result, and (c) divide by Δ. Of course to make these calculations feasible, the computer language implementing the calculations should support complex arithmetic and ideally have an automatic dispatching mechanism so that only one implementation of each function is required. In contrast to numerical integration of the joint system (1), the analytic method is embarrassingly parallelizable across parameters.

The following straightforward Julia routine for computing sensitivities

~~~
function differential(f::F, p, d) where F
  fvalue = real(f(p)) # function value
  df = zeros(length(fvalue), 0) # empty differential
  for j = 1:length(p)
    p[j] = p[j] + d * im # perturb parameter
    fj = f(p) # compute perturbed function value
    p[j] = complex(real(p[j]), 0.0) # reset parameter
    df = [df imag(fj) ./ d] # concatenate jth partial
  end
  return (fvalue, df) # return the differential
end
~~~

requires a function 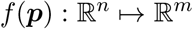 of a real vector ***p*** declared as complex. The perturbation scalar *d* should be small and real, say 10^−10^ to 10^−12^ in double precision. If the parameters *p_j_* vary widely in magnitude, then a good heuristic is to perturb *p_j_* by *p_j_di* instead of *di*. The returned value df is an *m* × *n* real matrix. The Julia commands `real` and `complex` effect conversions between real and complex numbers, and Julia substitutes im for 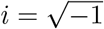. We will employ these functions later in some case studies.

A recent extension [13] of the analytic method facilitates accurate approximation of second derivatives. The relevant formula is

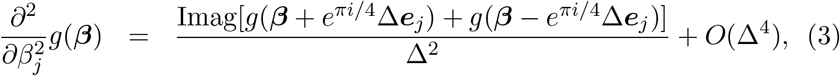

where 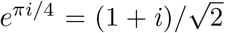. Roundoff errors can now occur, but are usually manageable. Recently we discovered how to extend the analytic method to approximate mixed partials. Our derivation is condensed into the following equations

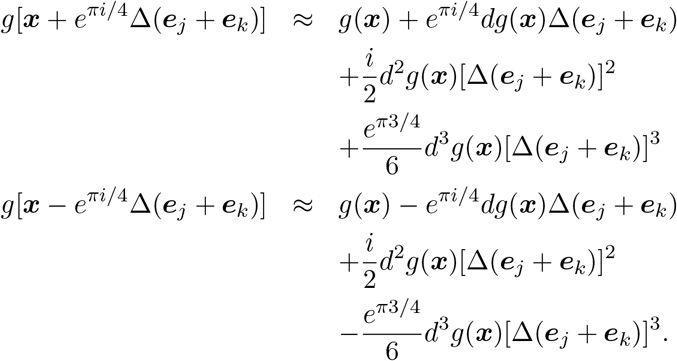

This approximation is accurate to order *O*(Δ^6^) and allows us to infer that

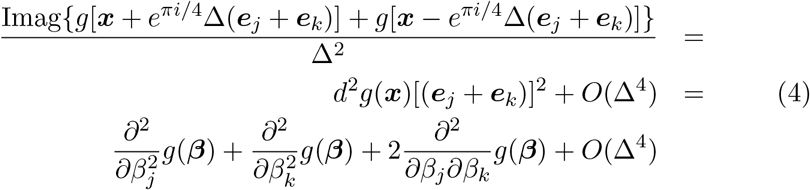

Since we can approximate 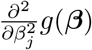 and 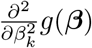, we can now approximate 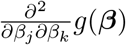 to order *O*(Δ^4^). These approximations are derived in the Appendix.

The Julia code for computing second derivatives

~~~
function hessian(f::F, p, d) where F
  d2f = zeros(length(p), length(p)) # hessian
  dp = d * (1.0 + 1.0 * im) / sqrt(2)
  for j = 1:length(p) # compute diagonal entries of d2f
    p[j] = p[j] + dp
    fplus = f(p)
    p[j] = p[j] - 2 * dp
    fminus = f(p)
    p[j] = complex(real(p[j]), 0.0) # reset parameter
    d2f[j, j] = imag(fplus + fminus) / d^2
  end
  for j = 2:length(p) # compute off diagonal entries
    for k = 1:(j - 1)
      (p[j], p[k]) = (p[j] + dp, p[k] + dp)
      fplus = f(p)
      (p[j], p[k]) = (p[j] - 2 * dp, p[k] - 2 * dp)
      fminus = f(p)
      (p[j], p[k]) = (complex(real(p[j]), 0.0), complex(real(p[k]), 0.0))
      d2f[j, k] = imag(fplus + fminus) / d^2
      d2f[j, k] = (d2f[j, k] - d2f[j, j] - d2f[k, k]) / 2
      d2f[k, j] = d2f[j, k]
    end
  end
  return d2f
end
~~~

operates on a scalar-valued function *f*(*u*) of a real vector ***p*** declared as complex. Because roundoff error is now a concern, the perturbation scalar *d* should be in the range 10^−3^ to 10^−6^ in double precision. The returned value *d*^2^*f* is a symmetric matrix.

Another technique one can use to calculate the derivatives of model solutions is to differentiate the numerical algorithm that calculates the solution. This can be done with computational tools collectively known as automatic differentiation [CITE]. Forward mode automatic differentiation is performed by carrying forward the Jacobian-vector product at each successive calculation. This is accomplished by defining higher-dimensional numbers, known as dual numbers [14], coupled to primitive functions *f*(***x***) through the action

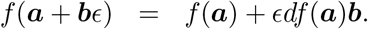

Here *ϵ* is a dimensional marker, similar to the complex *i*, which is a two-dimensional number. For a composite function *f* = *f*_2_ ∘ *f*_1_, the chain rule is *df*(***a***)***b*** = *df*_2_[*f*_1_(***a***)][*df*_1_(***a***)***b***]. The *i*th column of the Jacobian appears in the expression *f*(***x***+***e**_i_ϵ*) = *f*(***x***)+*ϵ*∇*_i_f*(***x***). Since computational algorithms can be interpreted as the composition of simpler functions, one need only define the algorithm on a small set of base cases (such as +, *, sin, and so forth, known as the primitives) and then apply the accepted rules in sequence to differentiate more elaborate functions. The ForwardDiff.jl package in Julia does this by defining dispatches for such primitives on a dual number type and provides convenience functions for easily extracting common objects like gradients, Jacobians, and Hessians. Hessians are calculated by layering automatic differentiation twice on the same algorithm to effectively take the derivative of a derivative.

In this form, forward mode automatic differentiation shares many similarities to the analytic methods described above without the requirement that the extension of *f*(***x***) be complex analytic. Of course, at every stage of the calculation *f*(***x***) must be differentiable. The pros and cons of this weaker but still restrictive assumption will be discussed in the results section. Conveniently, automatic differentiation allows for arbitrarily many derivatives to be calculated simultaneously. By defining higher-dimensional dual numbers that act independently via 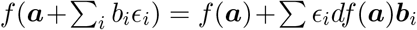, one can calculate entire Jacobians in a single function call 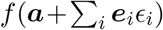. This use of higher-dimensional dual numbers is a practice known as chunking. Chunking reduces the number of primal (non-derivative) calculations required for computing the Jacobian. Because the ForwardDiff.jl package uses chunking by default, we will investigate the extent to which this detail is applicable in biological models.

## 3 Case Studies

We now explore applications of differential sensitivity to a few core models in oncology and epidemiology.

### 3.1 CARRGO Model

The CARRGO model [15] was designed to capture the tumor-immune dynamics of CAR T-cell therapy in glioma. The CARRGO model generalizes to other tumor cell-immune cell interactions. Its governing system of ODEs

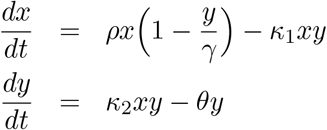

follows cancer cells *x* as prey and CAR T-Cells *y* as predators. This model captures Lotka-Volterra dynamics with logistic growth of the cancer cells. Our numerical experiments assume the parameter values and initial conditions

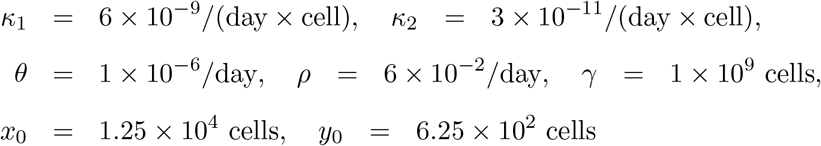

suggested by Sahoo et al. [15].

A traditional sensitivity analysis hinges on solving the system of ODEs for a variety of parameters and displaying the solution at a chosen future time across an interval or rectangle of parameter values. Figure 1 shows how *x*(*t*) and *y*(*t*) vary at *t* = 1000 days under joint changes of *κ*_1_ and *κ*_2_, where *κ*_1_ is the rate at which cancer cells are destroyed in an interaction with an immune cell, and *κ*_2_ is the rate at which immune cells are recruited after such an interaction. This type of analysis directly portrays how a change in one or two parameters impacts the outcome of the system. As might be expected, the number of cancer cells *x*(*t*) strongly depends on *κ*_1_ but weakly on *κ*_2_. In contrast, the number of immune cells *y*(*t*) depends comparably on both parameters, perhaps due to the fact that the initial population of immune cells is much smaller than the initial population of cancer cells.

**Figure 1:**
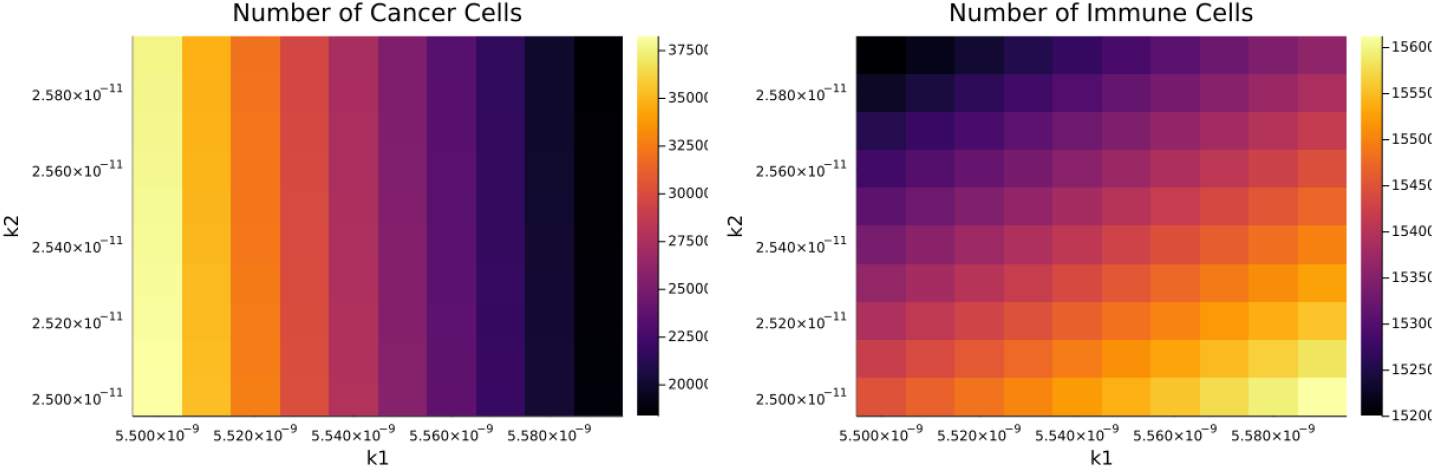
Sensitivity of Cancer Cells in the CARRGO Model

There are limitations to this type of sensitivity analysis. How many solution curves should be examined? What time is most informative in displaying system changes? Is it really necessary to compute sensitivity over such a large range of parameters when the trends are so clear? These ambiguities cloud our understanding and require far more computing than is necessary. Differential sensitivity successfully addresses these concerns. Gradients of solutions immediately yield approximate solutions in a neighborhood of postulated parameter values. The relative importance of different parameters in determining species levels is obvious from inspection of the gradient. Furthermore, modern software easily delivers the gradient along entire solution trajectories. There is no need to solve for an entire bundle of neighboring solutions.

Differential assessment is far more efficient. The required calculations involve solving an expanded system of ordinary differential equations just once under the partial derivative method or solving the system once for each parameter under the analytic method. Either way, the differential method is much less computationally intensive than the traditional method of solving the ODE system over an interval for each parameter or over a rectangle for each pair of parameters. Here is our brief Julia code for computing sensitivity via the analytic method.

~~~
using DifferentialEquations, Plots
function sensitivity(x0, p, d, tspan)
  problem = ODEProblem{true}(ODE, x0, tspan, p)
  sol = solve(problem, saveat = 1.0) # solve ODE
  (lp, ls, lx) = (length(p), length(sol), length(x0))
  solution = Dict{Int, Any}(i => zeros(ls, lp + 1) for i in 1:lx)
  for j = 1:lx # record solution for each species
    @views solution[j][:, 1] = sol[j, :]
  end
  for j = 1:lp
    p[j] = p[j] + d * im # perturb parameter
    problem = ODEProblem{true}(ODE, x0, tspan, p)
    sol = solve(problem, saveat = 1.0) # resolve ODE
    p[j] = complex(real(p[j]), 0.0) # reset parameter
    @views sol .= imag(sol) / d # compute partial
    for k = 1:lx # record partial for each species
      @views solution[k][:,j + 1] = sol[k, :]
    end
  end
  return solution
end
function ODE(dx, x, p, t) # CARRGO model
  dx[1] = p[4] * x[1] * (1 - x[1] / p[5]) - p[1] * x[1] * x[2]
  dx[2] = p[2] * x[1] * x[2] - p[3] * x[2]
end
p = complex([6.0e-9, 3.0e-11, 1.0e-6, 6.0e-2, 1.0e9]); # parameters
x0 = complex([1.25e4, 6.25e2]); # initial values
(d, tspan) = (1.0e-13, (0.0, 1000.0)); # step size and time interval in days
solution = sensitivity(x0, p, d, tspan); # find solution and partials
CARRGO1 = plot(solution[1][:, 1], label = “x1”, xlabel= “days”,
ylabel = “cancer cells x1”, xlims = (tspan[1],tspan[2]))
CARRGO2 = plot(solution[1][:, 2], label = “d1×1”, xlabel= “days”,
ylabel = “p1 sensitivity”, xlims = (tspan[1],tspan[2]))
~~~

In the Julia code the parameters *κ*_1_, *κ*_2_, *θ*, *ρ*, and *γ* and the variables *x* and *y* exist as components of the vector ***p*** and ***x***, respectively. The two plot commands construct solution curves for cancer and its sensitivity to perturbations of *κ*_1_.

Figure 2 reinforces the conclusions drawn from the heatmaps, but more clearly and quantitatively. In brief, the scaled sensitivity of cancer cells *x*(*t*) is much more dependent on carrying capacity *γ* than on birth rate *ρ*, and even more so than on the destruction rate *κ*_1_. Interestingly, the number of immune cells *y*(*t*) is much more sensitive to the recruitment rate *κ*_2_ than to the natural death rate *θ*.

**Figure 2:**
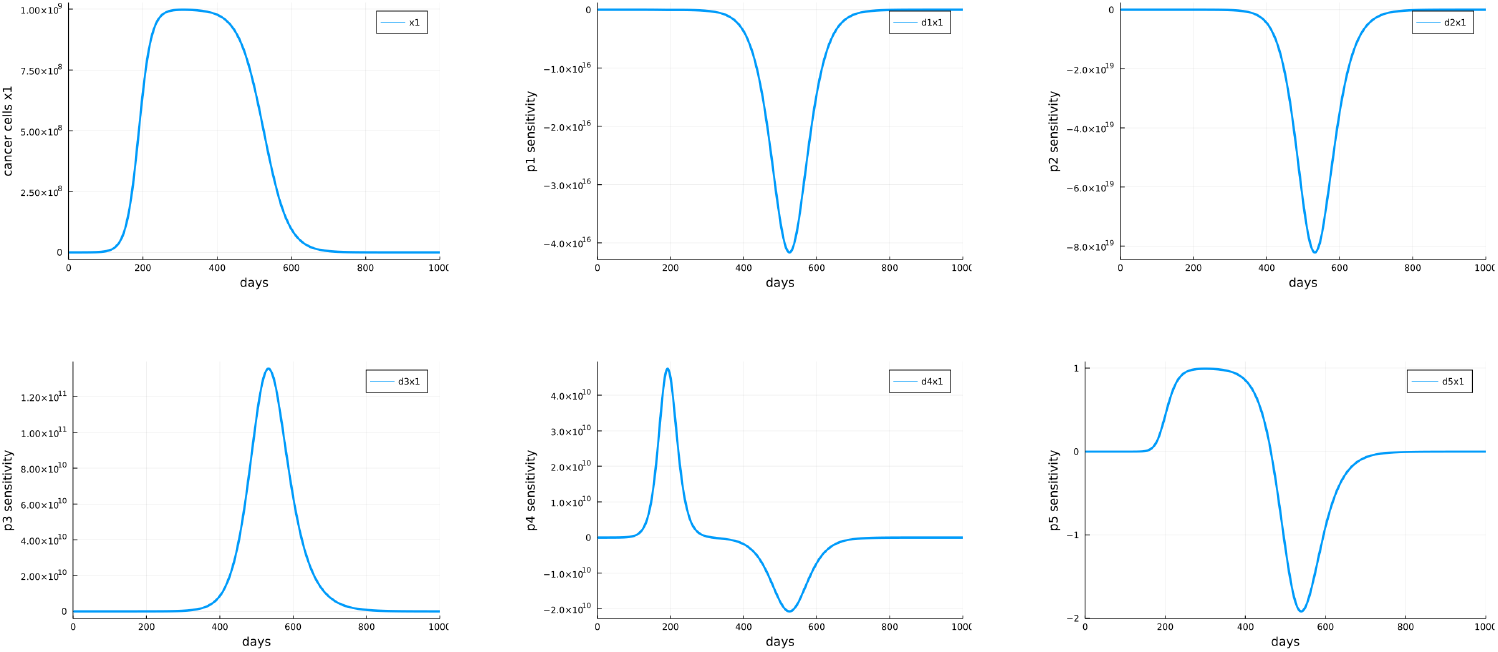
Sensitivity of Cancer Cells in the CARRGO Model

### 3.2 Deterministic SIR Model

The deterministic SIR model follows the number of infectives *I*(*t*), the number of susceptibles *S*(*t*), and the number of recovereds *R*(*t*) during the course of an epidemic. These three subpopulations satisfy the ODE system

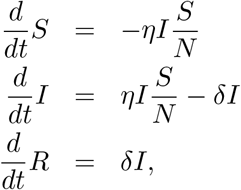

where *η* is the daily infection rate per encounter and *δ* is the daily rate of progression to immunity per person.

For SARS-CoV-2, current estimates [16] of *η* range from 0.0012 to 0.48, and estimates of *δ* range from 0.0417 to 0.0588 [17]. As an alternative to solving the extended set of differential equations, we again use the analytic method to evaluate parameter sensitivities.

The following Julia code for the analytic method incorporates the sensitivity function of the previous problem.

~~~
function ODE(dx, x, p, t) # Covid model
  N = 3.4e8 # US population size
  dx[1] = - p[1] * x[2] * x[1] / N
  dx[2] = p[1] * x[2] * x[1] / N - p[2] * x[2]
  dx[3] = p[2] * x[2]
end
p = complex([0.2, (0.0417 + 0.0588) / 2]); # parameters
x0 = complex([3.4e8, 100.0, 0.0]); # initial values
(d, tspan) = (1.0e-10, (0.0, 365.0)) # 365 days
solution = sensitivity(x0, p, d, tspan);
Covid = plot(solution[1][:, :], label = [“x1” “d1×1” “d2×1”],
        xlabel = “days”, xlims = (tspan[1],tspan[2]))
~~~

Our parameter choices roughly capture the American experience. Figure 3 plots the susceptible curve and its sensitivities. In this case all three curves conveniently occur on comparable scales.

**Figure 3:**
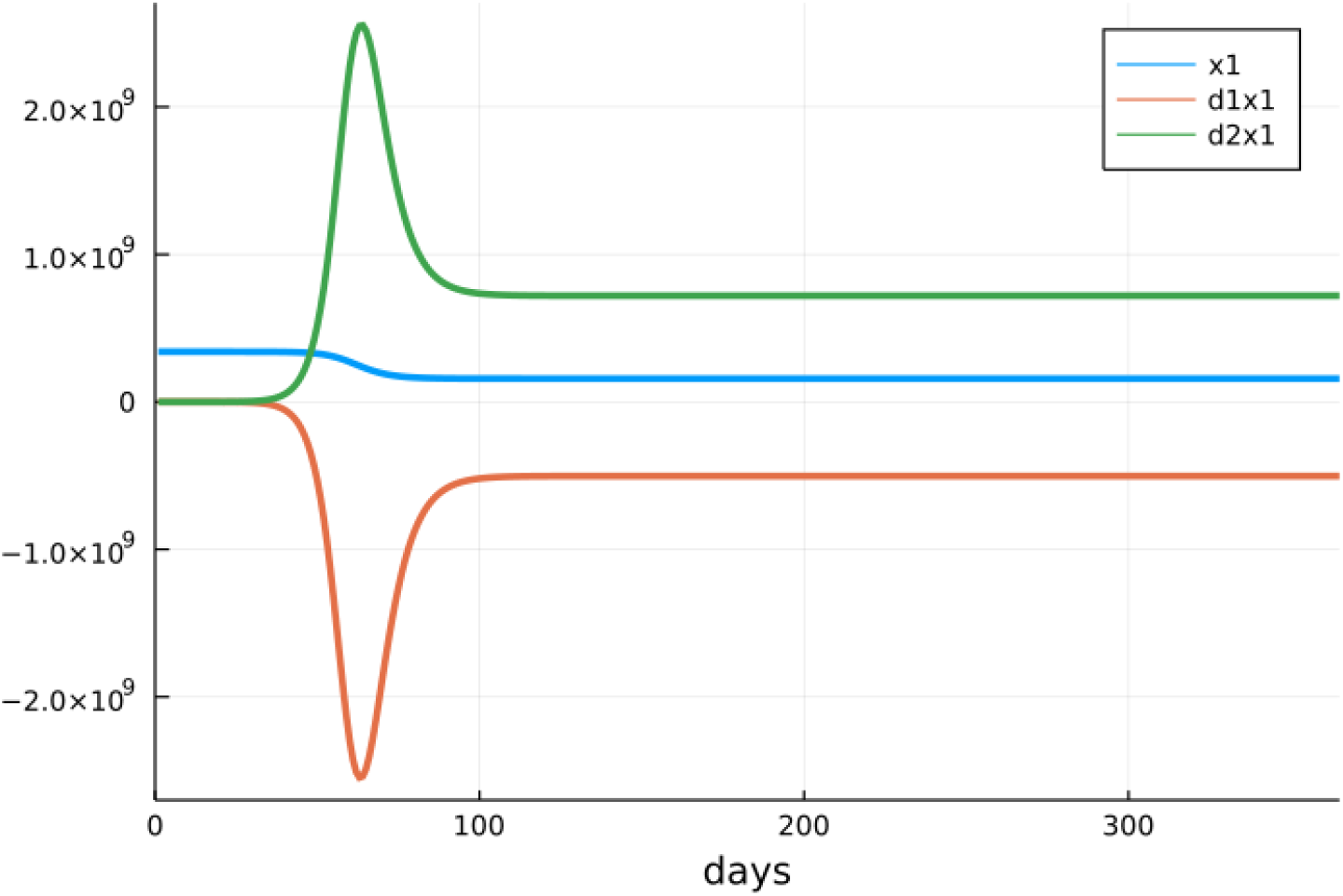
Sensitivities of Susceptibles in the Covid Model

### 3.3 Second-Order Expansions of Solution Trajectories

In predicting nearby solution trajectories, the second-order Taylor expansion

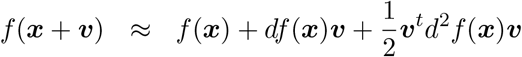

obviously improves accuracy over the first-order expansion

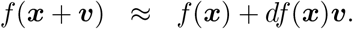

Figure 4 displays predicted trajectories for the SIR model when all parameters *p_i_* are replaced by *p_i_*(1 + *U_i_*), where each *U_i_* is chosen uniformly from (−0.25, 0.25). The figure vividly confirms the improvement in accuracy in passing from a first-order to a second-order approximation. The more non-linear the solution trajectory becomes, the more improvement is evident.

**Figure 4:**
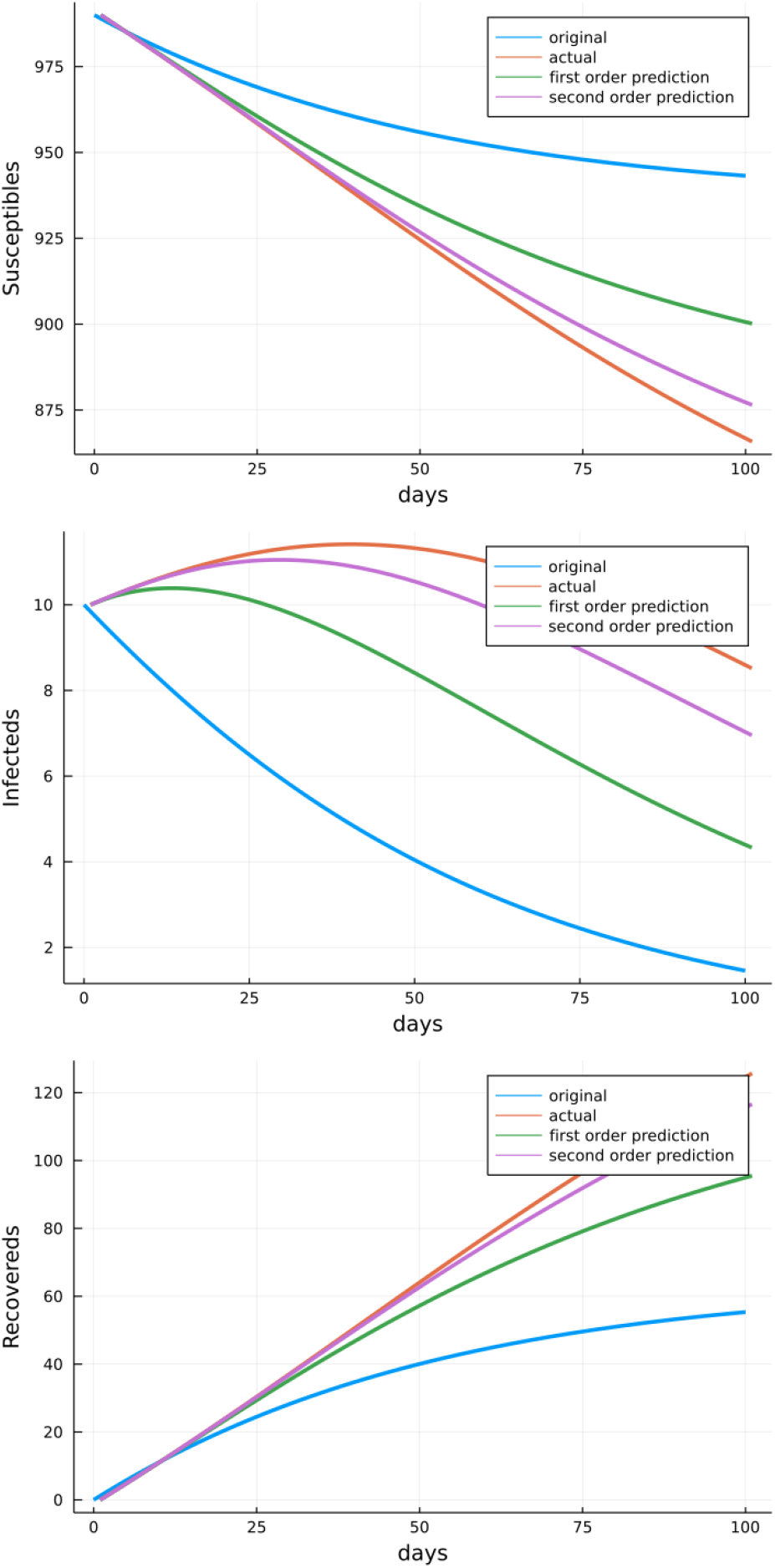
Model Trajectories for SIR Model Calculated Using First and Second Differentials

For example, the middle panel of Figure 4 shows that the solution trajectory of infected individuals bends dramatically with a change in parameters. This behavior is much better reflected in the second-order prediction compared to the first-order prediction. The Euclidean distance between the actual and predicted trajectories at the sampled time points is about 30.0 in the first-order case and only about 9.82 in the second-order case, a reduction of over 65% in prediction error. By contrast, the trajectory of the recovered individuals steadily increases in a nearly linear fashion. The bottom panel of Figure 4 shows that the first-order prediction now remains reasonably accurate over a substantial period of time. Even so, the discrepancy between the predicted solutions grows so that by day 100 the absolute difference between the first-order prediction and the actual trajectory exceeds 140, compared to about 39.0 for the second-order prediction. Thus, calculating second-order sensitivity is helpful in both highly non-linear systems and systems with long time scales.

### 3.4 Stochastic SIR Model

We now illustrate sensitivity calculations in the stochastic SIR model. This model postulates an original population of size *n* with *i* infectives and *s* susceptibles. The parameters *δ* and *η* again capture the rate of progression to immunity and the infection rate per encounter. Since extinction of the infectives is certain, it makes sense to focus on the time to elimination of the infectives. It is also convenient to follow the vector (*i, n*), where *n* = *i* + *s* is the sum of the number of infectives *i* plus the number of susceptibles *s*. Elementary arguments now show that the mean time *t_in_* to elimination of all infectives satisfies the recurrence

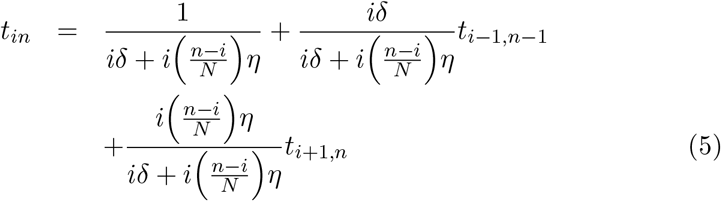

for 0 < *i* < *n* together with the boundary conditions

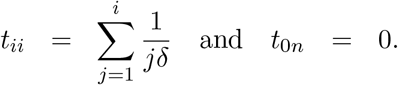

The expression for *t_ii_* stems from adding the expected time for the *i* → *i* – 1 transition, plus the expected time *i* – 1 → *i* – 2, and so forth. This system of equations can be solved recursively for *i* = *n*, *n* – 1, … 0 starting with *n* = 1. Once the values for a given *n* are available, *n* can be incremented, and a new round is initiated. Ultimately the target size *n* = *N* is reached. Taking partial derivatives of the recurrence (5) yields a new system of recurrences that can also be solved recursively in tandem with the original recurrence. The analytic method is easier to implement and comparable in accuracy to the partial derivative method.

Another important index of the SIR process is the mean number of infectives *m_in_* ever generated starting with *i* initial infectives and *n* total people. These expectations can be calculated via the recurrences

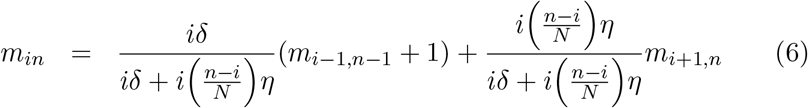

for 0 < *i* < *n* together with the boundary conditions

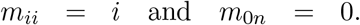

Obviously, one can compute the sensitivities of the *m_in_* to parameter perturbations in almost exactly the same way as the *t_in_*. Here is the Julia code for the two means and their sensitivities via the analytic method. Note how our earlier differential function plays a key role.

~~~
function SIRMeans(p)
  (delta, eta) = (p[1], p[2])
  M = Matrix(zeros(eltype(p), N + 1, N + 1)) # mean matrix
  T = similar(M) # time to extinction matrix
  for n = 1:N # recurrence relations loop
    for j = 0:(n - 1)
      i=n-j
      a = i * delta # immunity rate
      if i == n # initial conditions
        M[i + 1, n + 1] = i
        T[i + 1, n + 1] = T[i, i] + 1 / a
      else
        b = i * (n - i) * eta / N # infection rate
        c = 1 / (a + b)
        M[i + 1, n+1] = a * c * (M[i, n] + 1) + b * c * M[i + 2, n + 1]
        T[i + 1, n+1] = c *(1 + a * T[i, n] + b * T[i + 2, n + 1])
      end
    end
  end
  return [M[:, N + 1]; T[:, N + 1]]
end
p = complex([0.2, (0.0417 + 0.0588) / 2]); # delta and beta
(N, d) = (100, 1.0e-10);
@time (f, df) = differential(SIRMeanTime, p, d);
~~~

The left column of Figure 5 displays the expected total number of individuals infected (top) and the expected number of days to extinction (bottom) for the stochastic SIR model based on the parameters mentioned in Section 3.2. The middle column of Figure 5 depicts the derivatives of these quantities with respect to *η* (the infection rate), and the right column depicts the derivatives with respect to *δ* (the recovery rate).

**Figure 5:**
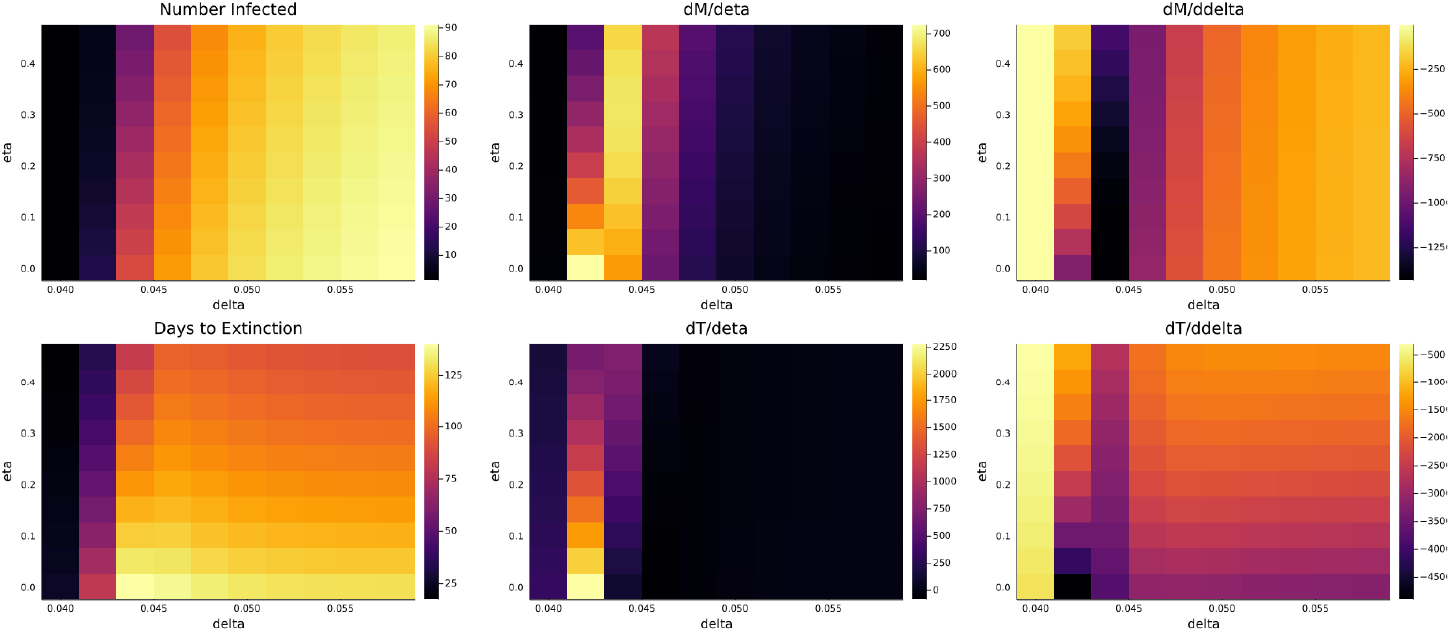
Sensitivity of Stochastic SIR Model

It is interesting to compare our method’s results to estimates from stochastic simulations. To see the difference in accuracy, we calculated the average number of individuals infected and the average time to extinction by stochastic simulation using the software package Biosimulator.jl [18]. Table 1 records the analytic and simulated means of these outcomes under the parameter values pertinent to Figure 3. As Table 1 indicates, the simulated means over *r* = 100 runs are roughly comparable to the analytic means, but the standard errors of the simulated means are large. Because the standard errors decrease as 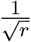, it is difficult to achieve much accuracy by simulation alone. In more complicated models, simulation is so computationally intensive and time consuming that it is nearly impossible to achieve accurate results. Of course, the analytic method is predicated on the existence of an exact solution or an algorithm for computing the same.

**Table 1:**
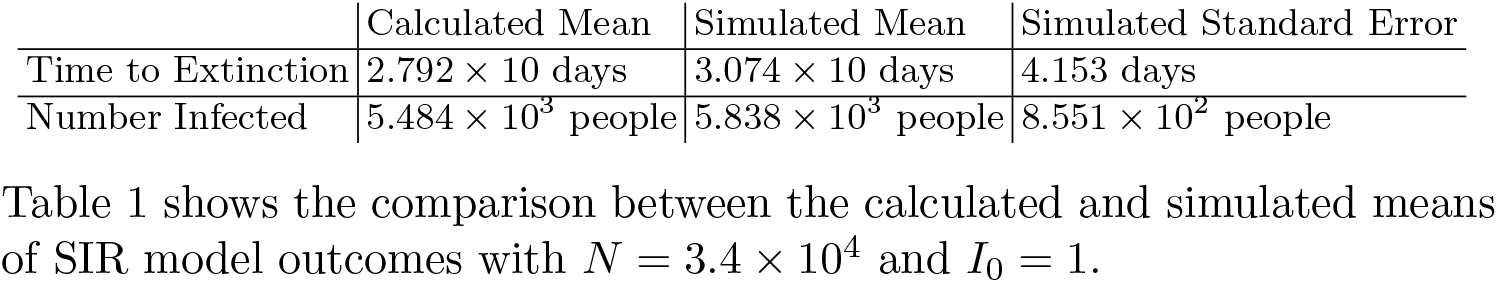
SIR Model Outcomes

|  | Calculated Mean | Simulated Mean | Simulated Standard Error |
| --- | --- | --- | --- |
| Time to Extinction | $2.792 \times 10$ days | $3.074 \times 10$ days | 4.153 days |
| Number Infected | $5.484 \times 10^3$ people | $5.838 \times 10^3$ people | $8.551 \times 10^2$ people |

Table 1 shows the comparison between the calculated and simulated means of SIR model outcomes with $N = 3.4 \times 10^4$ and $I_0 = 1$ .

Parameter sensitivities inform our judgment in interesting and helpful ways. For example, derivatives of both the total number of infecteds and the time to extinction with respect to *η* are very small except in a narrow window of the *δ* parameter. This suggests that we focus further simulations, sensitivity analysis, and possible interventions on the region of parameter space where *δ* is small and derivatives with respect to *η* tend to be large. Derivatives with respect to *δ* depend mostly on *η* except at very small values of *δ*. These conclusions are harder to draw from noisy simulations alone.

### 3.5 Branching Processes

Branching process models offer an opportunity for checking the accuracy of sensitivity calculations. For simplicity we focus on birth-death-migration processes [19]. These are multi-type continuous-time processes [9, 12] that can be used to model the early stages of an epidemic over a finite graph with *n* nodes, where nodes represent cities or countries. On node *i* we initiate a branching process with birth rate *β_i_* > 0 and death rate *δ*_i_ > 0. The migration rate from node *i* to node *j* is λ*_ij_* ≥ 0. All rates are per person, and each person is labeled by a node. Let 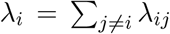 be the sum of the migration rates emanating from node *i*. Given this notation, the mean infinitesimal generator of the process is the matrix

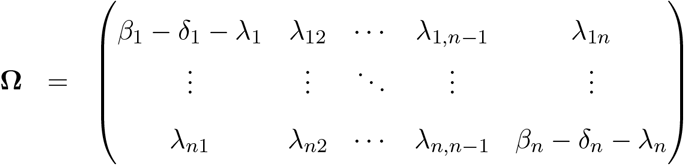

The entries of the matrix *e*^*t***Ω**^ = [*m_ij_*(*t*)] represent the expected number of people at node *j* at time *t* starting from a single person of type *i* at time 0. The process is irreducible when the pure migration process corresponding to choice *β_i_* = *δ_i_* = 0 for all *i* is irreducible. Henceforth, we assume the process is irreducible and let **Γ** denote the mean infinitesimal generator of the pure migration process. The process is subcritical, critical, or supercritical depending on whether the dominant eigenvalue *ρ* of **Ω** is negative, zero, or positive.

To determine the local sensitivity of *ρ* to a parameter *θ* [9, 20], suppose its left and right eigenvectors ***v*** and ***w*** are normalized so that ***vw*** = 1. Differentiating the identity **Ω*w*** = *ρ**w*** with respect to *θ* yields

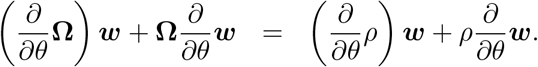

If we multiply this by ***v*** on the left and invoke the identities ***v*Ω** = *ρ**v*** and ***vw*** = 1 we find that

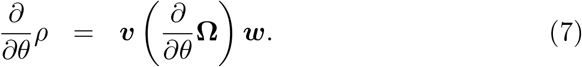

Because 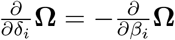, it follows that an increase in *δ_i_* has the same impact on *ρ* as the same decrease in *β_i_*. The sensitivity of ***v*** and ***w*** can be determined by an extension of this reasoning [22]. The extinction probabilities *e_i_* of the birth-death-migration satisfy the system of algebraic equations

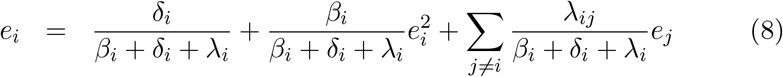

for all *i*. This is a special case of the vector extinction equation

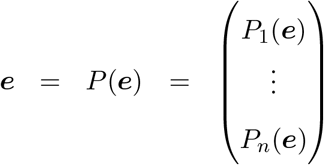

for a general branching process with offspring generating function *P_i_*(***x***) for a type *i* person [21]. For a subcritical or critical process, ***e*** = **1**. For a supercritical process all *e_i_* ∈ (0, 1). Iteration is the simplest way to find ***e***. Starting from ***e***_0_ = **0**, the vector sequence ***e**_n_* = *P*(***e***_*n*–1_) satisfies

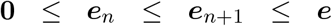

and converges to a solution of the extinction equations. Here all inequalities apply component-wise.

To find the differential [22] of the extinction vector ***e*** with respect to a vector ***θ*** of parameters, we assume that the branching process is supercritical and resort to implicit differentiation of the equation ***e***(***θ***) = *P*[***e***(***θ***), ***θ***]. The chain rule gives

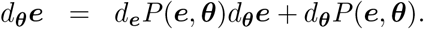

This equation has the obvious solution

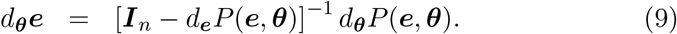

The indicated inverse does, in fact, exist, but the proof is a detour. Alternatively, one can compute an entire extinction curve ***e***(*t*) whose component *e_i_*(*t*) supplies the probability of extinction before time *t* starting from a single person of type *i* [12]. This task reduces to solving a ODE 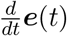 by the methods previously discussed. We leave the details to interested readers.

The following Julia code computes the sensitivities of the extinction probability for a two-node process by the analytic method.

~~~
using LinearAlgebra
function extinction(p)
  types = Int(sqrt(1 + length(p)) - 1) # length(p) = 2 * types + types^2
  (x, y) = (zeros(Complex, types), zeros(Complex, types))
  for i = 1:500 # functional iteration
    y = P(x, p)
    if norm(x - y) < 1.0e-16 break end
    x = copy(y)
  end
  return y
end
function P(x, p) # progeny generating function
  types = Int(sqrt(1 + length(p)) - 1) # length(p) = 2 * types + types^2
  delta = p[1: types]
  beta = p[types + 1: 2 * types]
  lambda = reshape(p[2 * types + 1:end], (types, types))
  y = similar(x)
  t = delta[1] + beta[1] + lambda[1, 2]
  y[1] = (delta[1] + beta[1] * x[1]^2 + lambda[1, 2] * x[2]) / t
  t = delta[2] + beta[2] + lambda[2, 1]
  y[2] = (delta[2] + beta[2] * x[2]^2 + lambda[2, 1] * x[1]) / t
  return y
end
delta = complex([1.0, 1.75]); # death rates
beta = complex([1.5, 1.5]); # birth rates
lambda = complex([0.0 0.5; 1.0 0.0]); # migration rates
p = [delta; beta; vec(lambda)]; # package parameter vector
(types, d) = (2, 1.0e-10)
@time (e, de) = differential(extinction, p, d)
~~~

To adapt the code to a different branching process model, one simply supplies the appropriate progeny generating function and necessary parameters.

The average number *a_ij_* of infected individuals of type *j* ultimately generated by a single initial infected individual of type *i* is also of interest. The matrix ***A*** = (*a_ij_*) of these expectations can be calculated via the matrix equation

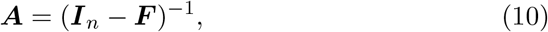

where ***F*** is the offspring matrix

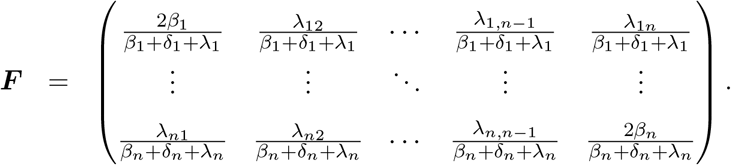

One can determine the local sensitivity of the expected numbers of total descendants by differentiating the equation ***A*** = (***I**_n_* – ***F***)^−1^. The result

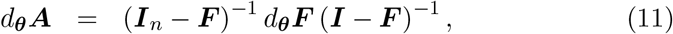

depends on the sensitivity of the expected offspring matrix ***F***. Julia code for the analytic method with two nodes follows.

~~~
function particles(p) # mean infected individuals generated
  types = Int(sqrt(1 + length(p)) - 1) # length(p) = 2 * types + types^2
  delta = p[1: types]
  beta = p[types + 1: 2 * types]
  lambda = reshape(p[2 * types + 1:end], (types, types))
  F = complex(zeros(types, types))
  t = delta[1] + beta[1] + lambda[1, 2]
  (F[1, 1], F[1, 2]) = (2 * beta[1] / t, lambda[1, 2] / t)
  t = delta[2] + beta[2] + lambda[2, 1]
  (F[2, 1], F[2, 2]) = (lambda[2, 1] / t, 2 * beta[2] / t)
  A = vec(inv(I - F)) # return as vector
end
delta = complex([1.0, 1.75]); # death rates
beta = complex([1.5, 1.5]); # birth rates
lambda = complex([0.0 0.5; 1.0 0.0]); # migration rates
p = [delta; beta; vec(lambda)]; # package parameter vector
(types, d) = (2, 1.0e-10)
@time (A, dA) = differential(particles, p, d)
~~~

## 4 Accuracy and Speed

We now assess the speed and accuracy of the computational schemes just described. In general, we measure accuracy by the Euclidean norms

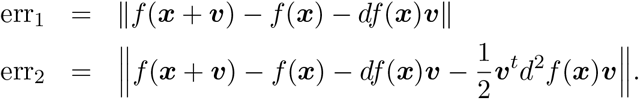

Other norms, such as the ℓ_1_ and ℓ_∞_ norms, yield similar results. In the ODE models, *f*(***x***) denotes a matrix trajectory so the Frobenius norm applies. This is scaled by 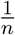, where *n* is the number of sampled time points. We also compare the analytic derivatives to the automatic derivatives delivered by DiffEqSensitivity.jl (abbreviated DES.jl) via normed differences. Finally, our speed comparisons offer a first look at the efficiency gains possible with multithreading. All computations were done in Julia version 1.6.2 on a Windows operating system with an Intel(R) Core (TM) i7-8565U CPU.

Table 2 records benchmarking results for three differential equations models. The top table shows the deterministic SIR model. This model, previously introduced in Section 3.2, serves as a straightforward benchmark for comparing the various differential sensitivity methods summarized earlier. For first-order approximations, we compare the analytic method with the automatic differentiation method available in DES.jl [3]. The table shows that the analytic method offers comparable accuracy but slower compute times than automatic differentiation. It is noteworthy that more naive implementations of forward mode differentiation do not take advantage of internal parallelization (chunking) and are consequently much slower than the analytic method. Results for forward mode differentiation with and without internal parallelization as implemented in ForwardDiff.jl [14] can be found in Appendix C.

**Table 2:** Time and Accuracy per Time Point for ODE System Derivatives over the Interval [0, t_end_]

| SIR Model |  |  |  |  |  |  |
| --- | --- | --- | --- | --- | --- | --- |
| | Time ($\mu s$) | | | Accuracy | | |
| | $t_{\text{end}} = 10$ | $t_{\text{end}} = 100$ | $t_{\text{end}} = 1000$ | $t_{\text{end}} = 10$ | $t_{\text{end}} = 100$ | $t_{\text{end}} = 1000$ |
| Forward DES.jl $\nabla F$ | $8.02 \times 10^1$ | $2.30 \times 10^2$ | $1.77 \times 10^3$ | 1.19 | $2.75 \times 10^2$ | $9.86 \times 10^3$ |
| Analytic $\nabla F$ | $2.44 \times 10^2$ | $9.94 \times 10^2$ | $8.82 \times 10^3$ | 1.19 | $2.75 \times 10^2$ | $9.86 \times 10^3$ |
| Multi-Threaded Analytic $\nabla F$ | $2.49 \times 10^2$ | $1.03 \times 10^3$ | $1.22 \times 10^4$ | 1.19 | $2.75 \times 10^2$ | $9.86 \times 10^3$ |
| Parallel FD.jl $\nabla(\nabla F)$ | $2.07 \times 10^2$ | $4.28 \times 10^2$ | $3.92 \times 10^3$ | $1.15 \times 10^{-1}$ | $7.24 \times 10^1$ | $6.78 \times 10^3$ |
| Multi-Threaded Parallel FD.jl $\nabla(\nabla F)$ | $1.71 \times 10^2$ | $4.29 \times 10^2$ | $3.95 \times 10^3$ | $1.15 \times 10^{-1}$ | $7.24 \times 10^1$ | $6.78 \times 10^3$ |
| Analytic $D^2 F$ | $8.86 \times 10^2$ | $3.76 \times 10^3$ | $5.20 \times 10^4$ | $1.15 \times 10^{-1}$ | $7.24 \times 10^1$ | $6.78 \times 10^3$ |
| Multi-Threaded Analytic $D^2 F$ | $8.77 \times 10^2$ | $3.26 \times 10^3$ | $5.57 \times 10^4$ | $1.15 \times 10^{-1}$ | $7.24 \times 10^1$ | $6.78 \times 10^3$ |

| CARRGO Model |  |  |  |  |  |  |
| --- | --- | --- | --- | --- | --- | --- |
| | Time ($\mu s$) | | | Accuracy | | |
| | $t_{\text{end}} = 10$ | $t_{\text{end}} = 100$ | $t_{\text{end}} = 1000$ | $t_{\text{end}} = 10$ | $t_{\text{end}} = 100$ | $t_{\text{end}} = 1000$ |
| Forward DES.jl $\nabla F$ | $1.21 \times 10^2$ | $3.69 \times 10^2$ | $2.30 \times 10^3$ | 6.19 | $2.80 \times 10^5$ | $1.46 \times 10^8$ |
| Analytic $\nabla F$ | $4.41 \times 10^2$ | $2.14 \times 10^3$ | $1.76 \times 10^4$ | 6.19 | $2.80 \times 10^5$ | $1.46 \times 10^8$ |
| Multi-Threaded Analytic $\nabla F$ | $4.43 \times 10^2$ | $2.17 \times 10^3$ | $2.58 \times 10^4$ | 6.19 | $2.80 \times 10^5$ | $1.46 \times 10^8$ |
| Parallel FD.jl $\nabla(\nabla F)$ | $1.23 \times 10^3$ | $4.48 \times 10^3$ | $4.38 \times 10^4$ | $1.13 \times 10^{-1}$ | $5.01 \times 10^4$ | $4.23 \times 10^7$ |
| Multi-Threaded Parallel FD.jl $\nabla(\nabla F)$ | $1.20 \times 10^3$ | $4.60 \times 10^3$ | $4.66 \times 10^4$ | $1.13 \times 10^{-1}$ | $5.01 \times 10^4$ | $4.23 \times 10^7$ |
| Analytic $D^2 F$ | $2.38 \times 10^3$ | $1.43 \times 10^4$ | $1.22 \times 10^5$ | $1.12 \times 10^{-1}$ | $4.96 \times 10^4$ | $7.93 \times 10^7$ |
| Multi-Thread Analytic $D^2 F$ | $2.00 \times 10^3$ | $1.14 \times 10^4$ | $1.25 \times 10^5$ | $1.12 \times 10^{-1}$ | $4.96 \times 10^4$ | $7.93 \times 10^7$ |

| MCC Model |  |  |  |  |  |  |
| --- | --- | --- | --- | --- | --- | --- |
| | Time ($\mu s$) | | | Accuracy | | |
| | $t_{\text{end}} = 10$ | $t_{\text{end}} = 100$ | $t_{\text{end}} = 1000$ | $t_{\text{end}} = 10$ | $t_{\text{end}} = 100$ | $t_{\text{end}} = 1000$ |
| Forward DES.jl $\nabla F$ | $6.12 \times 10^2$ | $9.35 \times 10^3$ | $2.51 \times 10^4$ | $3.47 \times 10^{-3}$ | $7.56 \times 10^{-3}$ | $1.54 \times 10^{-1}$ |
| Analytic $\nabla F$ | $2.71 \times 10^3$ | $1.51 \times 10^4$ | $1.08 \times 10^5$ | $3.47 \times 10^{-3}$ | $7.56 \times 10^{-3}$ | $1.54 \times 10^{-1}$ |
| Multi-Threaded Analytic $\nabla F$ | $2.56 \times 10^3$ | $1.51 \times 10^4$ | $1.08 \times 10^5$ | $3.47 \times 10^{-3}$ | $7.56 \times 10^{-3}$ | $1.54 \times 10^{-1}$ |
| Parallel FD.jl $\nabla(\nabla F)$ | $1.84 \times 10^4$ | $7.44 \times 10^4$ | $2.04 \times 10^5$ | $1.26 \times 10^{-3}$ | $2.07 \times 10^{-3}$ | $4.21 \times 10^{-2}$ |
| Multi-Threaded FD.jl $\nabla(\nabla F)$ | $1.76 \times 10^4$ | $7.77 \times 10^4$ | $2.04 \times 10^5$ | $1.26 \times 10^{-3}$ | $2.07 \times 10^{-3}$ | $4.21 \times 10^{-2}$ |
| Analytic $D^2 F$ | $4.78 \times 10^4$ | $2.81 \times 10^5$ | $1.00 \times 10^6$ | $1.27 \times 10^{-3}$ | $1.92 \times 10^{-3}$ | $3.89 \times 10^{-2}$ |
| Multi-Threaded Analytic $D^2 F$ | $2.56 \times 10^4$ | $1.74 \times 10^5$ | $1.36 \times 10^6$ | $1.27 \times 10^{-3}$ | $1.92 \times 10^{-3}$ | $3.89 \times 10^{-2}$ |

As we expect, second-order approximations are more accurate in prediction. For second-order methods, we compare the double Jacobian method found in ForwardDiff.jl with the second-order analytic method. In contrast to the analytic method, forward differentiation slows dramatically in calculating the Hessian directly, and only speeds up if we instead calculate the gradient of the gradient. The ForwardDiff.jl package currently does not offer this method natively, so the analytic method provides a simple way to improve the speed of computing second derivatives. While both of these methods allow for multi-threading, this model is too low dimensional to show much of an improvement.

The middle subtable in Table 2 records benchmarking results for the CARRGO model previously described in Section 3.1. This model serves as an example of a stiff system of differential equations. We can immediately see the impact of stiffness in the loss of accuracy at time points 100 and 1000. Stiffness highlights the added value of the second-order approximations. With stiff ODEs, high quality integrators are a must. In the analytic method, one must also choose the perturbation interval *d* carefully in computing second derivatives. Careful calibration of *d* improves the accuracy of the CARRGO second derivative analytic prediction to the level of automatic differentiation.

To explore whether parallelization improves computational speed in high-dimensional models, we now turn a third model summarized in the bottom subtable of Table 2. This ODE model of the mammalian cell cycle (MCC) was constructed by Gerard and Goldbetor [23] and is explained in Appendix B. The model comprises 11 equations and 15 parameters and captures aspects of cell reproduction and cycling mediated by chemical signaling from cell-state dependent proteins such as through tumor repressors, transcription factors, and other DNA replication checkpoints. The model relies on cell state as opposed to cell mass and nicely replicates sequential progression along the cell cycle. The model clearly demonstrates the increase in speed from parallelization of both the analytic method and forward differentiation.

We now turn to the stochastic SIR model and compare our two suggested methods for computing the derivatives of the mean number of infected individuals (*M*) and the mean time to extinction (*T*). Both methods depend on the recurrences (5) and (6). The direct differentiation method relies on differentiating these recurrences, while the analytic method simply perturbs parameter values and recomputes solutions. Table 3 records the speed and accuracy of the two methods. As expected, computation speed varies quadratically with the number of individuals in the system *N*. The analytic method proves to be twice as fast as the direct method.

**Table 3:**
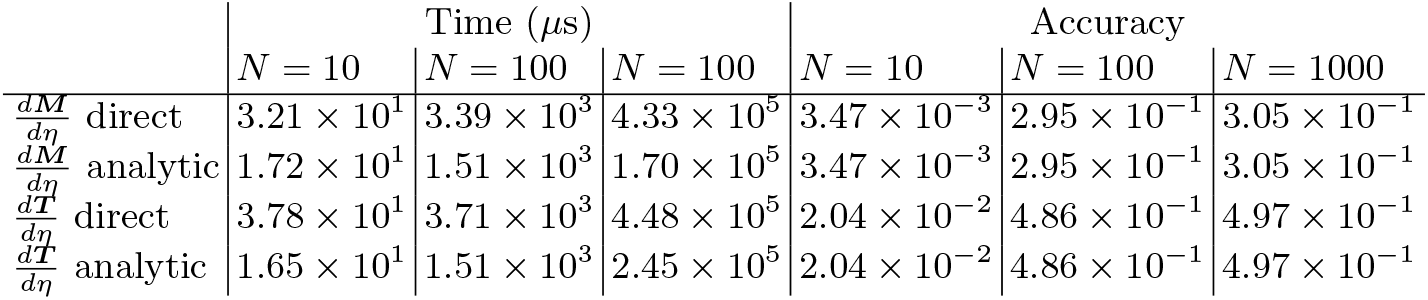
Time and Accuracy in the Stochastic SIR Model

| | Time ($\mu s$) | | | Accuracy | | |
| --- | --- | --- | --- | --- | --- | --- |
| | $N = 10$ | $N = 100$ | $N = 1000$ | $N = 10$ | $N = 100$ | $N = 1000$ |
| $\frac{dM}{d\eta}$ direct | $3.21 \times 10^1$ | $3.39 \times 10^3$ | $4.33 \times 10^5$ | $3.47 \times 10^{-3}$ | $2.95 \times 10^{-1}$ | $3.05 \times 10^{-1}$ |
| $\frac{dM}{d\eta}$ analytic | $1.72 \times 10^1$ | $1.51 \times 10^3$ | $1.70 \times 10^5$ | $3.47 \times 10^{-3}$ | $2.95 \times 10^{-1}$ | $3.05 \times 10^{-1}$ |
| $\frac{dT}{d\eta}$ direct | $3.78 \times 10^1$ | $3.71 \times 10^3$ | $4.48 \times 10^5$ | $2.04 \times 10^{-2}$ | $4.86 \times 10^{-1}$ | $4.97 \times 10^{-1}$ |
| $\frac{dT}{d\eta}$ analytic | $1.65 \times 10^1$ | $1.51 \times 10^3$ | $2.45 \times 10^5$ | $2.04 \times 10^{-2}$ | $4.86 \times 10^{-1}$ | $4.97 \times 10^{-1}$ |

The branching process model for the spread of COVID-19 is similar to the SIR model. Table 4 compares the direct method of calculating 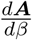 based on equation (11) to the analytic method based on equation (10). It also compares the direct method of calculating 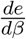 based on differentiating the recurrence (8) to the analytic method based on the same recurrence. In this case, the analytic method proves to be just as accurate as the direct methods, albeit slower in some cases. Other evidence not shown suggests that the analytic method can reliably evaluate sensitivities where solutions depend on linear algebra and/or recurrence relations. Unless derivatives are quite complicated, direct methods will generally be faster. In computing second derivatives, we expect the tables will be turned.

**Table 4:** Time and Accuracy in the Branching Process Model

| | Time ($\mu s$) | | | Accuracy | | |
| --- | --- | --- | --- | --- | --- | --- |
| | $N = 10$ | $N = 100$ | $N = 100$ | $N = 10$ | $N = 100$ | $N = 1000$ |
| $\frac{dA}{d\beta}$ direct | 4.28 | $2.46 \times 10^3$ | $1.09 \times 10^5$ | $1.02 \times 10^{-2}$ | $7.41 \times 10^{-4}$ | $1.43 \times 10^{-5}$ |
| $\frac{dA}{d\beta}$ analytic | $1.46 \times 10^2$ | $4.28 \times 10^3$ | $1.68 \times 10^5$ | $1.02 \times 10^{-2}$ | $7.41 \times 10^{-4}$ | $1.43 \times 10^{-5}$ |
| $\frac{de}{d\beta}$ direct | $1.29 \times 10^3$ | $8.29 \times 10^4$ | $7.75 \times 10^6$ | $9.23 \times 10^{-5}$ | $9.25 \times 10^{-7}$ | $1.85 \times 10^{-15}$ |
| $\frac{de}{d\beta}$ analytic | $9.69 \times 10^2$ | $5.64 \times 10^4$ | $1.20 \times 10^7$ | $9.23 \times 10^{-5}$ | $9.25 \times 10^{-7}$ | $1.85 \times 10^{-15}$ |

## 5 Discussion

Our purpose throughout has been to demonstrate the ease and utility of incorporating differential sensitivity analysis in dynamical modeling. Because models are always approximate, and parameters are measured imprecisely, uncertainty plagues all of dynamical modeling. Improving models is incremental and domain specific. Sensitivity analysis provides a handle on local parameter uncertainty. Assessing global parameter sensitivity is more challenging. It can be attacked by techniques such as Latin square hypercube sampling or Sobel quasi-random sampling, but these become infeasible in high dimensions [24]. Given the availability of appropriate software, differential sensitivity is computationally feasible, even for high dimensional systems.

In the case of stochastic models, traditional methods require costly and inaccurate simulation over a bundle of parameter values. Differential sensitivity is often out of the question. This limits the ability of researchers to understand a biological system and how it responds to parameter changes. If a system index such as a mean, variance, extinction probability, or extinction time can be computed by a reasonable algorithm, then differential parameter sensitivity analysis can be undertaken. We have indicated in a handful of examples how this can be accomplished.

In summary, across many models representative of computational biology, we have found the following results to hold:

a. Chunked forward mode automatic differentiation and forward mode sensitivity analysis tend to be the most efficient on the tested models.
b. The analytic methods described in this manuscript are competitive and often outperform the unchunked version of forward mode automatic differentiation.
c. Shared memory multithreading of the analytic and forward mode automatic differentiation methods provides a performance gain in high-dimensional systems.
d. Adjoint methods tend to have better scalability to very large systems with more than 100 ODEs [25]. However, on our test problems the adjoint-based methods were not competitive.
e. Forward mode automatic differentiation method requires that each step of a calculation is differentiatiable. This renders it unusable for calculating the derivative of ensemble means of discrete state models, such as birth-death processes. For these cases, the analytic method outperforms direct numerical differentiation.

These conclusions are tentative, but supported by our limited number of biological case studies.

We note that the performance differences may change depending on the efficiency of the implementations. The Julia DifferentialEquations.jl library and its DiffEqSensitivity.jl package have been shown to be highly efficient, outperforming other libraries in both equation solving and derivative calculations in Python, MATLAB, C, and Fortran [25, 26] (Details on the current state of performance can be found at https://github.com/SciML/SciMLBenchmarks.jl.) The automatic differentiation implementations in machine learning libraries optimize array operations much more than scalar operations. This can work to the detriment of forward mode AD. Similarly, forward mode AD sensitivity analysis is more amendable to MATLAB or Python style vectorization that improves performance by reducing interpreter overhead. Therefore, our conclusions can serve as guidelines for the case where all implementations are well-optimized. However, when using programming languages with high overheads or without compile-time optimization of the automatic differentiation passes, the balance in efficiency shifts more favorably towards the analytic method.

One last point worth making is on the coding effort required by the various methods. Both automatic differentiation and the analytic method have comparable accuracy when applied to systems of ODEs, with automatic differentiation having the advantage in speed when it is implemented with an additional level of parallelization. However, the analytic method can easily be generalized to other kinds of objective functions and may be more straightforward to implement for those less sophisticated in computer science. While automatic differentiation is the basis of many large scientific packages, the code required for the analytic methods is fully contained within this manuscript and is easily transferable to other programming languages with similar dispatching on complex numbers. This hard to measure benefit should not be ignored by practicing biologists who simply wish to quickly arrive at reasonably fast code.

## Acknowledgments

We wish to thank Chris Elrod for assistance in adding multithreading to parts of our software. We wish to thank Janet Sinsheimer, Mary Sehl, and Xiang Ji for helpful comments on the manuscript and biological applications.

## Appendix A: Derivation of Second Derivative Analytic Method

To prove the formulas for approximating partial derivatives stated in the text, we first note that any analytic function *f*(***z***) of several variables can be expanded in a locally convergent power series about every point ***z*** of its open domain of definition. If we choose a real direction vector ***v***, then the function *g*(*w*) = *f*(***z*** + *w**v***) is locally analytic in the complex plane 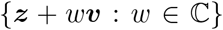 and can be expanded in a power series around *w* = 0. Thus,

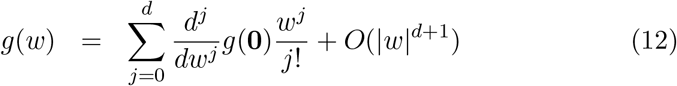

for any integer *d* ≥ 0. Now consider the setting where ***z*** has real components. If *f*(***z***) is real valued, then the derivatives 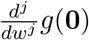 will be real as well. One can exploit this fact in approximating the derivatives. For example, if *w* = *i*, then *w^j^* rotates among the four values 1, *i*, −1, and –*i*. Because the terms of the expansion (12) alternate between real and imaginary values, the first partial derivative formula (2) holds. For the choice *w* = *e*^*πi*/4^, the powers *w^d^* rotate among the eight values 1, *e*^*πi*/4^, *i*, *ie*^*πi*/4^, –1, – *e*^*πi*/4^, –*i*, and –*ie*^*πi*/4^. The powers (–*w*)^*j*^ = (–1)*^j^w^j^* agree in this regard except for sign. Hence, the terms in the expansion of the sum

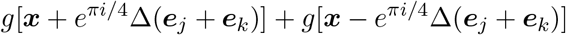

alternately cancel and reinforce. Thus, the first five terms of the expansion are real, 0, imaginary, 0, real, 0. It follows that the imaginary part of the sum is accurate to order *O*(Δ^6^) and that the approximations (3) and (4) are accurate to order *O*(Δ^4^).

## Appendix B: Mammalian Cell Cycle Model

The Mammalian Cell Cycle Model presented in [23] reduces to a system of ordinary differential equations representing the interaction of cyclin-dependent kinases (Cdk) with Cdk inhibitors, growth factors, and other proteins that regulate the development of mammalian cells. The model includes characteristics such as cell cycling, tumor repressor initiated progression control,and cell cycle completion. The ODE system representing the model is

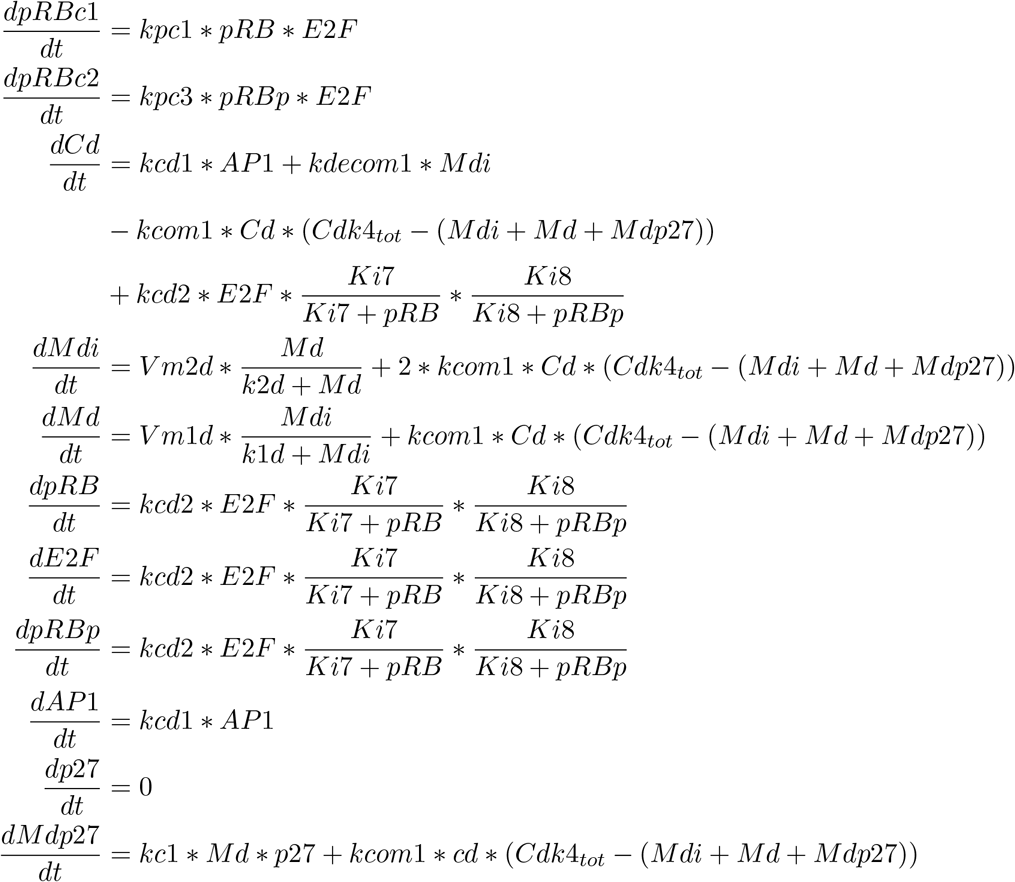

The initial values of each compartment and parameter are defined as

| Compartment | Initial Value ($\mu$ mol) | Parameter | Value |
| --- | --- | --- | --- |
| pRBc1 | 0.1 | kpc1 | 0.05 |
| pRBc2 | 0.05 | kpc3 | 0.025 |
| Cd | 0.01 | kcd1 | 0.4 |
| Mdi | 0.01 | Ki8 | 2.0 |
| Md | 0.01 | Ki7 | 0.1 |
| pRB | 0.0 | kcd2 | 0.005 |
| E2F | 0.0 | kdecom1 | 0.1 |
| pRBp | 0.0 | k2d | 0.1 |
| AP1 | 0.0 | Cdk4<sub>tot</sub> | 1.5 |
| p27 | 0.0 | kcom1 | 0.175 |
| Mdp27 | 0.0 | Vm2d | 0.2 |
| — | — | k1d | 0.1 |
| — | — | Vm1d | 1.0 |
| — | — | kc1 | 0.15 |

## Appendix C: Supplementary Benchmarking Data

In addition to the methods for conducting the differential sensitivity analysis for systems of ODEs displayed in Table 2 (Section 4), we conducted benchmarking tests on some variations of these methods as well. Tables 5 and 6 summarize these benchmarks for the SIR Model. Additional first-order methods include using the ForwardDiff.jl [14] package to calculate the Jacobian with and without chunking and with and without multi-threading, and using a constant value for *ϵ* = 1 × 10^−12^ instead of one scaled by the parameters. Additional second order methods include using DES.jl (both adjoint and non-adjoint) to calculate the Hessian, using the Forward-Diff.Hessian method, using the ForwardDiff.jacobian(ForwardDiff.jacobian) method without chunking (with and without multithreading), and using the constant value *ϵ* = 1 × 10^−6^ in the analytic method.

**Table 5:** Time and Accuracy per Time Point for ODE System Derivatives over the Interval [0, t_end_] - Additional First Order Methods

| SIR Model Time in $\mu s$ | | | | | |
| --- | --- | --- | --- | --- | --- |
| $t_{\text{end}}$ | Parallel FD.jl<br>Gradient | Parallel FD.jl<br>Multi-Threaded Gradient | FD.jl<br>Multi-Threaded Gradient | FD.jl<br>Gradient | Constant Analytic<br>Gradient |
| 10 | $1.40 \times 10^2$ | $1.47 \times 10^2$ | $3.41 \times 10^2$ | $3.59 \times 10^2$ | $3.19 \times 10^2$ |
| 100 | $4.33 \times 10^2$ | $4.39 \times 10^2$ | $1.10 \times 10^3$ | $1.12 \times 10^3$ | $1.40 \times 10^3$ |
| 1000 | $3.20 \times 10^3$ | $4.79 \times 10^3$ | $8.73 \times 10^3$ | $1.15 \times 10^4$ | $1.28 \times 10^4$ |

| SIR Model Accuracy |  |  |  |  |  |
| --- | --- | --- | --- | --- | --- |
| $t_{\text{end}}$ | Parallel FD.jl<br>Gradient | Parallel FD.jl<br>Multi-Threaded Gradient | FD.jl<br>Multi-Threaded Gradient | FD.jl<br>Gradient | Constant Analytic<br>Gradient |
| 10 | 1.19 | 1.19 | 1.19 | 1.19 | 1.19 |
| 100 | $2.75 \times 10^2$ | $2.75 \times 10^2$ | $2.75 \times 10^2$ | $2.75 \times 10^2$ | $2.75 \times 10^2$ |
| 1000 | $9.86 \times 10^3$ | $9.86 \times 10^3$ | $9.86 \times 10^3$ | $9.86 \times 10^3$ | $9.86 \times 10^3$ |

**Table 6:** Time (ns) and Accuracy per Time Point for ODE System Derivatives over the Interval [0, t_end_] - Additional Second Order Methods

| SIR Model Time in $\mu s$ | | | | | |
| --- | --- | --- | --- | --- | --- |
| $t_{\text{end}}$ | DES.jl Adjoint<br>Hessian | DES.jl Non-Adjoint<br>Hessian | FD.jl<br>Double Jacobian | FD.jl Multi-Threaded<br>Double Jacobian | Constant Analytic<br>Hessian |
| 10 | $3.72 \times 10^4$ | $1.18 \times 10^4$ | $1.00 \times 10^3$ | $9.81 \times 10^2$ | $1.16 \times 10^3$ |
| 100 | $4.23 \times 10^5$ | $1.30 \times 10^5$ | $3.27 \times 10^3$ | $3.19 \times 10^3$ | $5.28 \times 10^3$ |
| 1000 | $8.67 \times 10^6$ | $2.37 \times 10^6$ | $4.22 \times 10^4$ | $4.26 \times 10^4$ | $5.61 \times 10^4$ |

| SIR Model Accuracy |  |  |  |  |  |
| --- | --- | --- | --- | --- | --- |
| $t_{\text{end}}$ | DES.jl Adjoint<br>Hessian | DES.jl Non-Adjoint<br>Hessian | FD.jl<br>Double Jacobian | FD.jl Multi-Threaded<br>Double Jacobian | Constant Analytic<br>Hessian |
| 10 | $1.15 \times 10^{-1}$ | $1.15 \times 10^{-1}$ | $1.15 \times 10^{-1}$ | $1.15 \times 10^{-1}$ | $7.53 \times 10^{-1}$ |
| 100 | $7.24 \times 10^1$ | $7.24 \times 10^1$ | $7.24 \times 10^1$ | $7.24 \times 10^1$ | $1.33 \times 10^2$ |
| 1000 | $6.78 \times 10^3$ | $6.78 \times 10^3$ | $6.78 \times 10^3$ | $6.78 \times 10^3$ | $7.15 \times 10^3$ |

## Notes

### Competing Interest Statement

The authors have declared no competing interest.

